# A unifying principle for global greenness patterns and trends

**DOI:** 10.1101/2023.02.25.529932

**Authors:** Wenjia Cai, Ziqi Zhu, Sandy P. Harrison, Youngryel Ryu, Han Wang, Boya Zhou, Iain Colin Prentice

**Affiliations:** Georgina Mace Centre for the Living Planet, Department of Life Sciences, Imperial College London, UK; Department of Earth System Science, Ministry of Education Key Laboratory for Earth System Modelling, Institute for Global Change Studies, Tsinghua University, Beijing 100084, China; School of Archaeological and Environmental Sciences, University of Reading, UK; Department of Landscape Architecture and Rural Systems Engineering, Seoul National University, South Korea

## Abstract

Vegetation cover regulates the exchanges of energy, water and carbon between land and atmosphere. Remotely-sensed fractional absorbed photosynthetically active radiation (fAPAR), a land-surface greenness metric, depends on carbon allocation to foliage while also controlling photon flux for photosynthesis. Greenness is thus both a driver and an outcome of gross primary production (GPP). An equation with just two (globally) fitted parameters describes annual maximum fAPAR (fAPAR_max_) as the smaller of a water-limited value, transpiring a constant fraction of annual precipitation, and an energy-limited value, maximizing annual plant growth. This minimalist description reproduces global greenness patterns, and the consistent temporal trends among remote-sensing products, as accurately as the best-performing dynamic global vegetation models. Widely observed greening is attributed to the influence of rising carbon dioxide on the light- and water-use efficiencies of GPP, augmented by wetting in some dry regions and warming in high latitudes. Limited regions show browning, attributed to drying.

---

The exchanges of carbon, water and energy between terrestrial ecosystems and the atmosphere are regulated by vegetation cover, often quantified by leaf area index (LAI)^1^. Plants’ use of photosynthetically active radiation (PAR) for photosynthesis depends on LAI through its relationship to the fraction of PAR absorbed by leaves (fAPAR), which depends on canopy architecture^2^ but can be represented at its simplest by Beer’s law, fAPAR ≈ 1 – exp (–*k*. LAI), with a constant extinction coefficient (*k* ≈ 0.5)^3^. Transpiration, tightly coupled to photosynthesis through stomatal regulation, constitutes the largest part of total global land-surface evaporation and therefore makes a major contribution to the global hydrological cycle^4^. LAI also influences the partitioning of net surface radiation to latent versus sensible heat fluxes, which exerts a first-order control on the surface energy balance – a key determinant of local and regional climates.

Global products based on optical remote sensing, representing the seasonal and interannual time course of fAPAR, are widely used as inputs to land biosphere diagnostic models. Most of these models adopt a light use efficiency (LUE) formulation, whereby gross primary production (GPP) – that is, total photosynthesis per unit land area – over periods of a week or longer is proportional to absorbed PAR (the product of fAPAR and incident PAR)^5^. This approach can be justified theoretically as a consequence of photosynthetic acclimation, which translates the saturation response of photosynthesis to PAR – observed on sub-daily time scales – to a proportionality on time scales similar to the turnover time of Rubisco, the primary carboxylating enzyme^6^. Dynamic global vegetation models (DGVMs), which form the land-surface component of many contemporary Earth System models, independently predict LAI as a by-product of the partitioning of biomass production among leaves, stems and roots. However, this partitioning is one of the less studied aspects of ecosystem function, and the existing model formulations have not been extensively tested.

Here we explore an alternative approach: seeking first-order explanations for the emergence of spatial patterns and temporal trends of fAPAR in terms of simple eco-evolutionary optimality principles, which have been successful in explaining many features of plant and ecosystem behaviour^7^. Ground-based and remote-sensing data are used to test the hypothesis that the annual maximum fAPAR (fAPAR_max_) can be represented as the lesser of two quantities: a water-limited value determined by the principle of ‘ecohydrological equilibrium’, which states that a fixed fraction of antecedent annual precipitation is available to support photosynthesis^8^; and an energy-limited value, which maximizes plant growth by balancing the benefit of photosynthesis against a cost that is assumed to be proportional to LAI. Conceptually, this cost comprises the construction and maintenance costs of leaves, plus the additional carbon allocation required to keep the leaves supplied with water and nutrients. Our approach was first applied to the Tibetan Plateau^9^ and used to explain observed, divergent responses of energy- and water-limited vegetation to recent environmental changes. Ref. 59 paved the way for the present, global application.

Implementation of this hypothesis is facilitated by the use of the P model^6,10,11^, a universal, first-principles LUE model for GPP that has no free parameters and requires no plant functional type distinctions apart from the separation of C_3_ and C_4_ photosynthetic pathways (see Online Methods). The P model’s formulation as a LUE model results in a proportionality between fAPAR, GPP and transpiration^12^. Under water limitation, optimal fAPAR is taken to be the value that transpires a globally invariant fraction (to be estimated from data) of antecedent annual precipitation. This is an optimality hypothesis, in the sense that it assumes plants adapt their rooting strategy to compensate for different temporal distributions of precipitation. Under energy limitation, optimal fAPAR is taken as the value yielding the largest excess of annual GPP over a canopy cost, which is assumed to be proportional to LAI. Optimization of LAI is assumed to be independent of leaf mass-per-area (LMA), as co-existing plant species typically very substantially in LMA – yet those with higher LMA do not display lower LAI^13^. We use remotely-sensed fAPAR rather than LAI as the quantity for comparison with model results in order to avoid problems associated with the saturation of reflectances at high LAI values, where fAPAR approaches 1 (ref. 14). We focus on the annual maximum value of fAPAR per 0.5° grid cell and year (fAPAR_max_) as plants must allocate enough carbon to foliage to achieve this maximum, and specifically on the 95^th^ percentile of the distribution of fAPAR_max_ within each grid cell in order to minimize the effects of disturbance, succession and land management. We do not consider phenology here, which can however be separately predicted (given an annual maximum LAI) based on the seasonal time course of potential GPP^15^.

## Optimality criteria

For C_3_ plants, the optimality equations (see Online Methods for derivations) are:

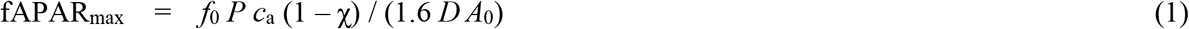

under water limitation and

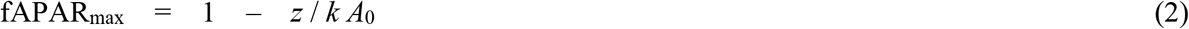

under energy limitation. Here fAPAR_max_ is the predicted annual maximum fAPAR. In equation (1), *f*_0_ is the fraction of annual precipitation (*P*, mol m^−2^ a^−1^) available for uptake by plants; *c*_a_ is the ambient partial pressure of CO_2_ (Pa); χ is the ratio of leaf-internal to ambient CO_2_; *D* is the vapour pressure deficit (Pa); and *A*_0_ is the potential GPP (mol m^−2^ a^−1^), which we define as the GPP predicted when fAPAR = 1 (i.e. the product of incident PAR and LUE, which is calculated by the P model). In equation (2), *z* is the foliage cost factor (mol m^−2^ a^−1^) and *k* is the extinction coefficient. The term χ is calculated by the P model as a function of growth temperature, vapour pressure deficit and atmospheric pressure, and LUE as a function of incident PAR, temperature and χ. Single, global values of the two free parameters (*f*_0_ and *z*) were estimated by non-linear regression (see Online Methods).

### Global comparisons to in situ data and spatial patterns

Ground-based measurements of green vegetation cover apply to small areas and can therefore be subject to large variation related to local soil conditions and disturbance history. Reasonable agreement was nonetheless obtained between a composite set of ground-based fAPAR^16,17^ and our predictions (Fig. 1). Mismatches were predominantly overpredictions, particularly in shrublands; the observed values are bounded above by a line close to the 1:1 line between observations and predictions. The geographic pattern of fAPAR_max_ predicted by this approach shows good agreement with the observed global pattern derived from MODIS data^18^ (Fig. 2).

**Figure 1:**
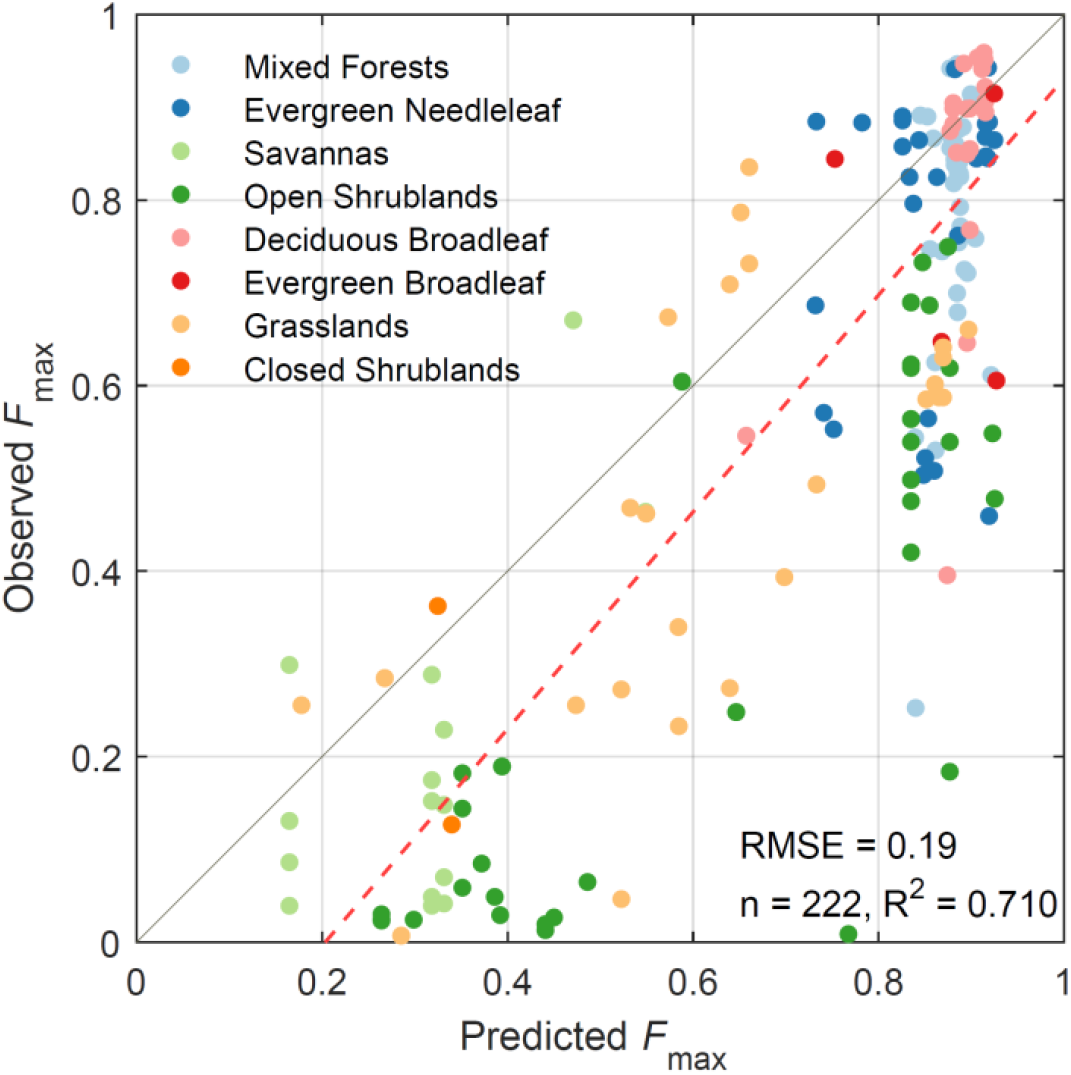
Comparison of predicted and in-situ observed annual maximum fAPAR at field sites. Observations are from the GBOV and OLIVE datasets. Predictions are from the theoretical model driven by environmental variables only; the model is incognizant of biome type. Biomes are shown with different colours. The dashed red ine is the ordinary least-squares regression line and the solid grey line is the 1:1 line. RMSE, root-mean-squared error of prediction; *n*, number of observations; *R*^2^, coefficient of determination.

**Figure 2:**
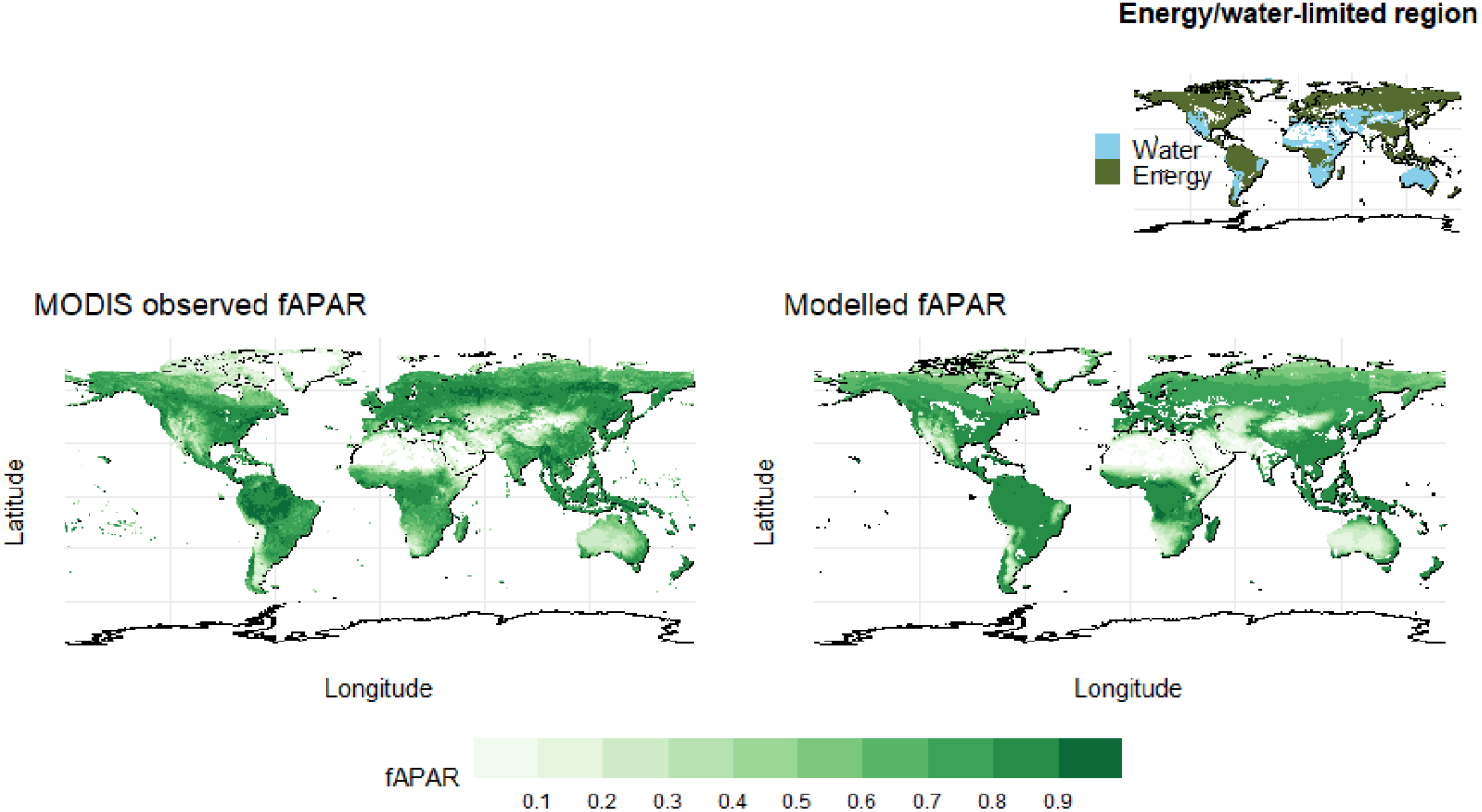
Predicted and observed (MODIS) annual maximum fAPAR. Energy- and water-limited regions, according to the model, are shown in the inset.

We compared the performance of our model with that of 15 DGVMs participating in the TRENDY project^19^ version 9 (Fig. 3). Our predictions performed as well as the best of the TRENDY models on the three criteria of *R*^2^, slope of the regression of observed versus modelled annual maximum fAPAR, and root-mean-squared error (RMSE) – achieving the highest *R*^2^ (0.95) tied with two other models (the model range is from 0.76 to 0.95), a regression slope within ± 0.02 of unity along with only four other models (the model range is from –0.18 to +0.13), and a low RMSE (0.15) equalled or marginally outperformed by only three other models (the model range is 0.14 to 0.33).

**Figure 3:**
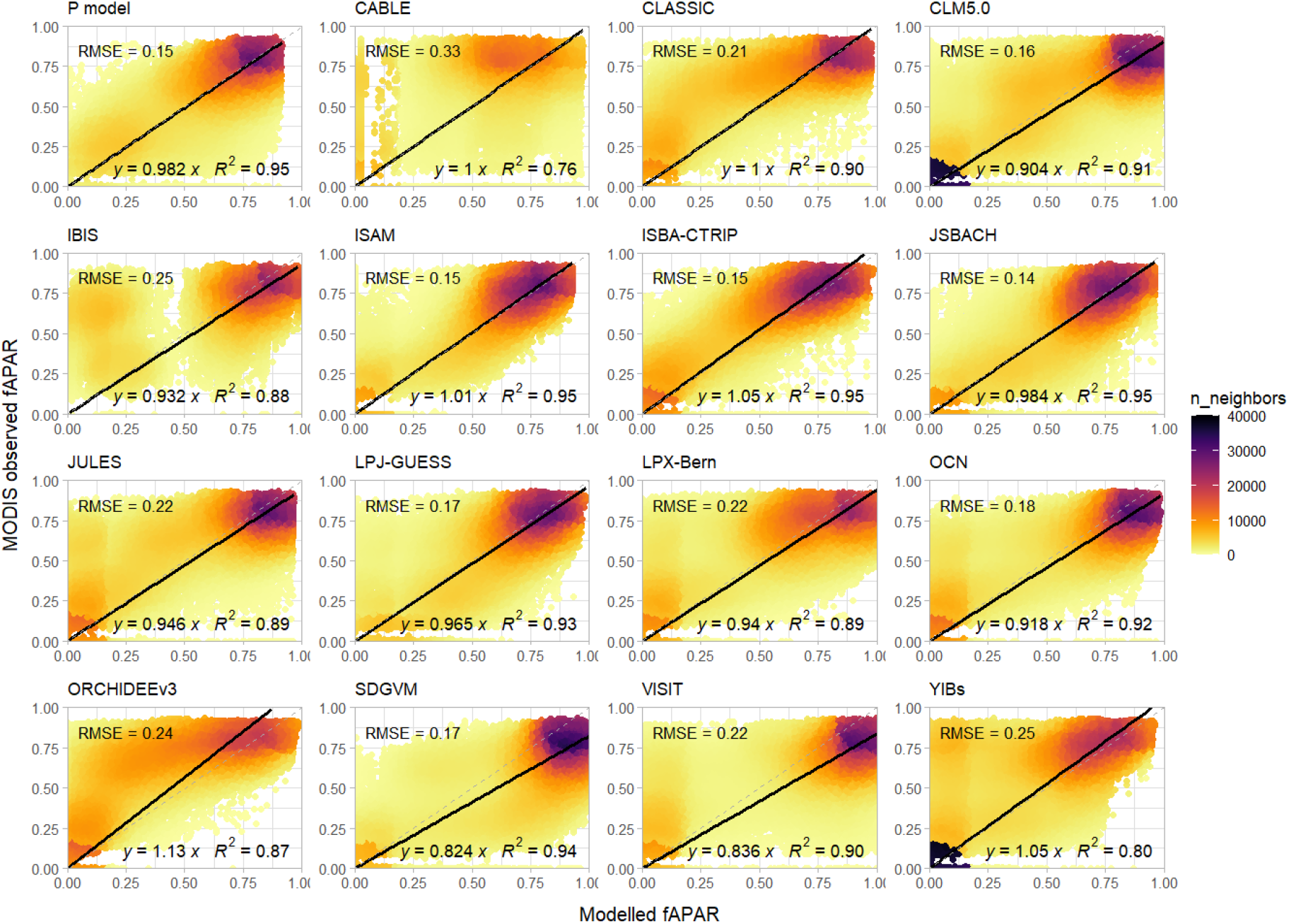
Comparison of simulated and observed (MODIS) annual maximum fAPAR. Simulated values are from this paper (P model) and the 15 models participating in the TRENDY project. Solid line is the ordinary least-squares regression line; grey dashed line is the 1:1 line. RMSE, root-mean-squared error of prediction; *y*, regression slope; *R*^2^, coefficient of determination.

### Trends during the MODIS era

Temporal trends in observed and modelled fAPAR_max_ during 2000–2017 were represented by non-parametric Mann-Kendall coefficients (see Online Methods). The evaluation of modelled temporal trends is hindered by relatively large, and generally unexplained, differences among remotely sensed products. Nonetheless, after eliminating grid cells where fitted trends showed opposite signs in different products, we can simulate the major features of observed trends (Fig. 4). Widespread greening was both observed and modelled. However, parts of semi-arid Central Asia and southwestern Africa, coastal California, the caatinga region of north-eastern Brazil and interior Argentina showed both observed and modelled browning trends. The model achieved a level of agreement in reproducing temporal trends comparable with that attained by the best-performing TRENDY models. *R*^2^ values greater than that obtained with the P model (0.37) were achieved by 40% of the TRENDY models. However, among these models, our model alone approached a regression slope of 1 (0.81) while others showed regression slopes of 0.63 or less. Across all models, *R*^2^ values ranged from 0.13 to 0.46, and regression slopes from 0.45 to 0.81. RMSE valued were similar (0.29 to 0.37) across all models.

**Figure 4:**
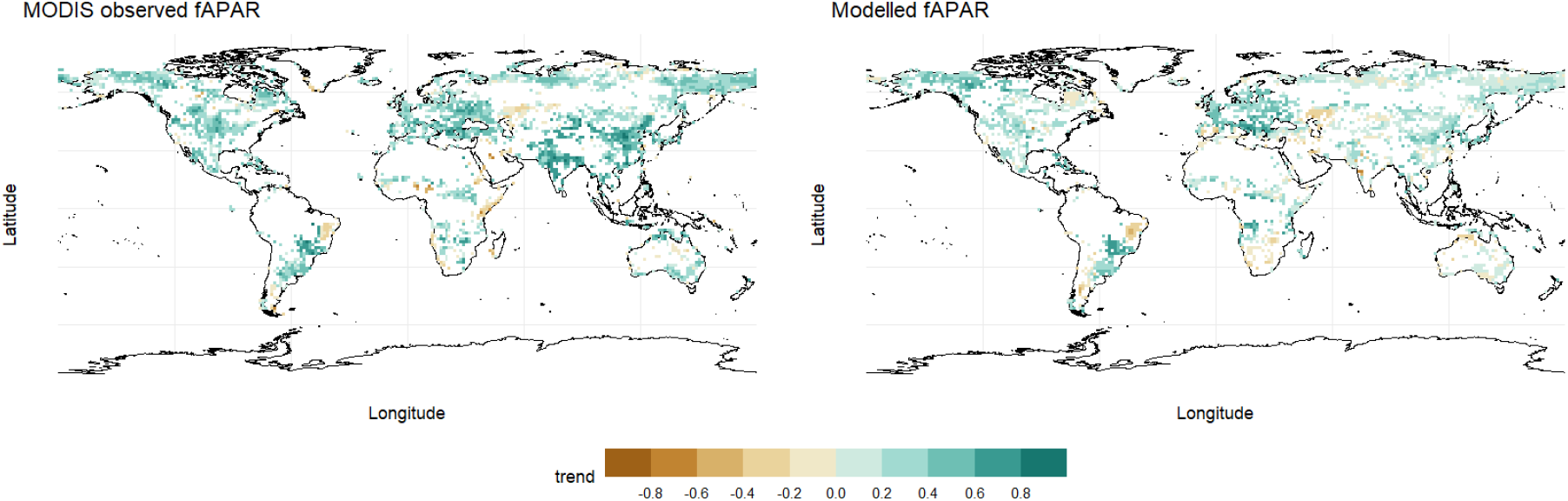
Predicted and observed (MODIS) trends in annual maximum fAPAR (Mann-Kendall coefficients). Values are not shown for grid cells with fAPAR > 0.85 in the initial year, or where different remotely-sensed products disagree on the sign of the trend.

**Figure 5:**
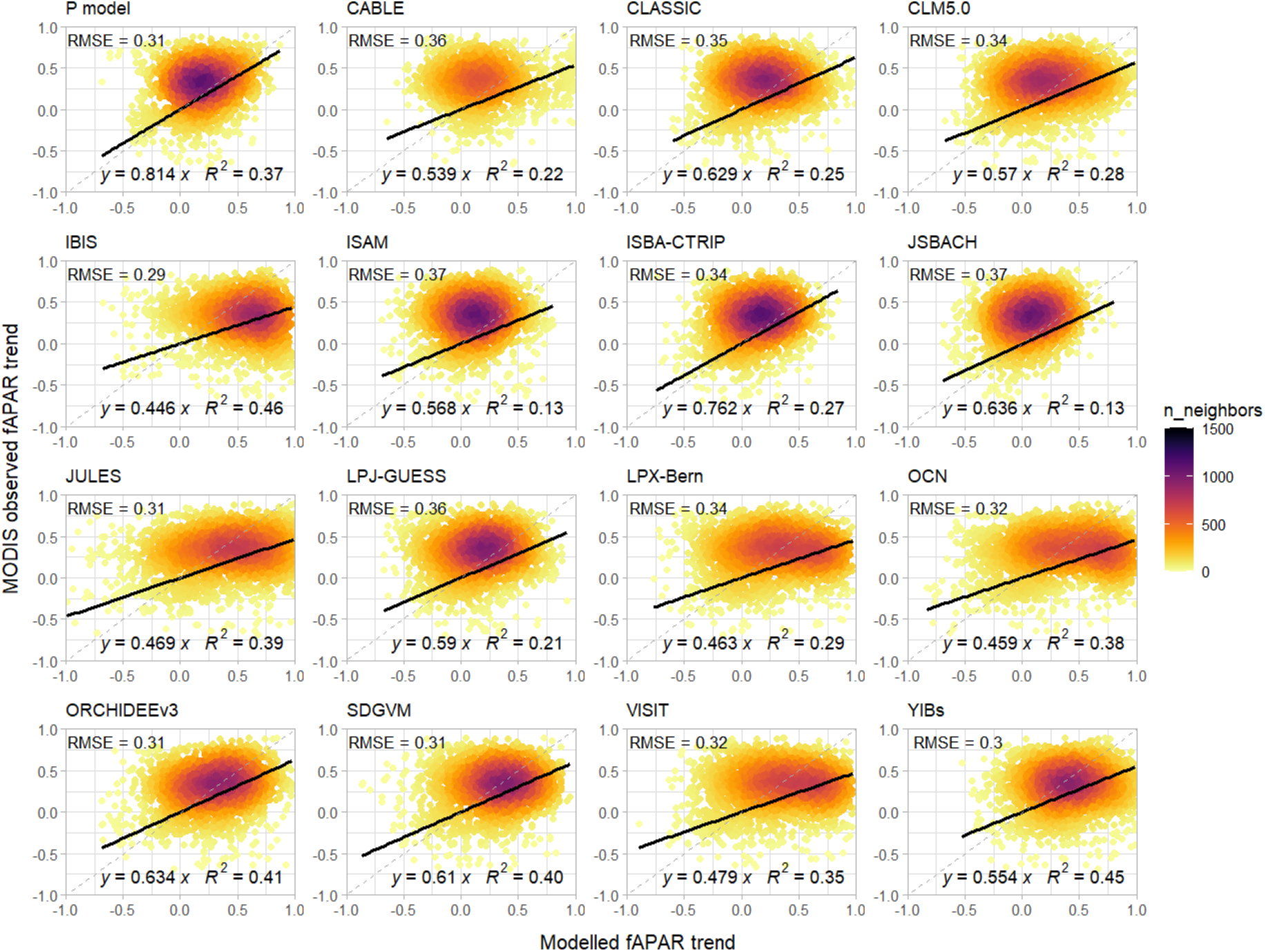
Comparison of simulated and observed (MODIS) trends in annual maximum fAPAR. Simulated values are from this paper (P model) and the 15 models participating in the TRENDY project. Solid line is the ordinary least squares regression line; grey dashed line is the 1:1 line. RMSE, root-mean-squared error of prediction; *y*, regression slope; *R*^2^, coefficient of determination.

**Figure 6:**
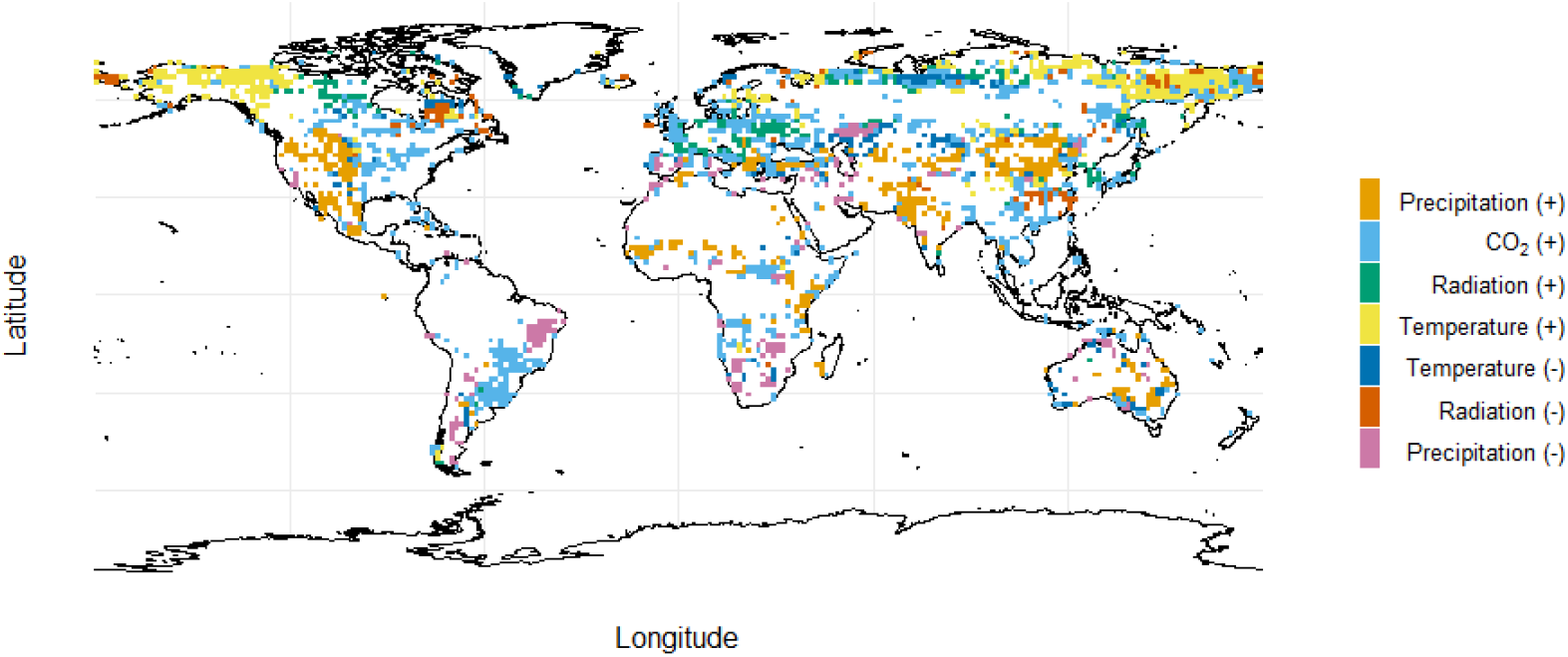
Dominant drivers of the trends in predicted annual maximum fAPAR, where (+) and (–) indicate a positive or negative effect of each driver. Values are not shown for grid cells with fAPAR > 0.85 in the initial year, or where different remotely-sensed products disagree on the sign of the trend.

For grid cells where most remotely sensed products agreed on the sign of the temporal trends, we estimated the dominant control of the trend by performing multiple simulations in which each environmental variable was held constant in turn in order to assess its influence (see Online Methods). Rising CO_2_ was implicated as the dominant cause of greening trends over much of the world, in agreement with previous analyses^20^. However, substantial areas showed greening trends that the model attributed to increasing precipitation, while smaller areas (including those listed previously) showed browning trends that the model attributed to declining precipitation. Increasing temperature is indicated to have led to greening in some regions of the northern high-latitudes, but to browning in a few other locations^20^. There are also regions, for example in parts of India and China, where observed greening has been stronger than predicted. This is likely due to direct human influence in the form of agricultural intensification and (in China) widespread reforestation^21^.

## Discussion

Our top-down approach intentionally glosses over potentially important details, such as the likelihood of environmental influences on *f*_0_ and *z*. Under water limitation, we have simply assumed that plants can access a fixed fraction of precipitation, adapting their root-zone capacity as needed to ensure this even in highly seasonal climates. Some theoretical work has pointed to an optimality constraint on this fraction, acting via the coordination of stomatal and hydraulic traits^22^. Yang et al.^23^ postulated the existence of a maximum ‘rain use efficiency’ and found, through an analogy with the Budyko curve, that high fAPAR sharpened the transition between energy and water limitation of GPP. Under energy limitation, we have assumed a fixed cost factor *z* for the maintenance and replenishment of the canopy. However, this factor is also likely to vary with environmental factors, including growth temperature, growing-season length and aridity^24^ as well as aspects of soil fertility, as less fertile soils are likely to induce higher costs in below-ground allocation for the uptake of nutrients^25^. Such dependencies would repay further study both from a theoretical standpoint, and in remotely sensed observations. Nonetheless, the fidelity of our minimalist model to the major spatial and temporal patterns in fAPAR data is notable, as is its ability to reproduce both aspects as well as or better than far more complex and comprehensive DGVMs.

Ecohydrological optimality concepts have a long but chequered history. Some specific hypotheses earlier proposed by Peter Eagleson (a pioneer of global ecohydrology) are now considered implausible from an eco-evolutionary perspective^26,27^. Here we have intentionally kept the formulations as simple as possible, to illustrate general principles, and to facilitate comparison with previous studies. By linking remotely sensed data with theory in this way, we hope that this work will contribute to the development of more comprehensive theory for the interactions of vegetation and its growth environment.

This research also has potential utility in providing one key component of a general optimality-based model for ecosystem function. Current global land models contain many *ad hoc* and untested ‘legacy’ elements – not least for the determination of carbon allocation to roots, shoots and leaves, for which there are few generally accepted principles. If the tendency of fAPAR can be predicted (and the costs of LAI quantified in terms of required below-ground allocation), it should be possible also to predict annual carbon profit – and thereby formulate an evolutionarily stable strategy for height competition among plants^28^. These elements are all required, if a new generation of land ecosystem models is to rest on secure foundations.

## Methods

We used the P model^6,10,11^ to provide the components of equations (1) and (2). The P model is based on the Farquhar–von Caemmerer–Berry photosynthesis model, but has the mathematical form of a LUE model. It simulates terrestrial ecosystem gross primary production (GPP) as a function of atmospheric CO_2_ concentration, air temperature, atmospheric pressure, vapour pressure deficit, incident photosynthetic photon flux density (PPFD), the fraction of incident PPFD absorbed by vegetation (fAPAR), root-zone soil moisture (θ) and C_4_ vegetation fraction. A detailed description can be found in ref. 10, where it is also shows that the model can account for observed trends in GPP at multiple eddy-covariance flux sites. Soil water stress was implemented following ref. 29, whereby χ is down-regulated under water stress following an empirical relationship with soil moisture content. A dynamic C_4_ vegetation fraction was simulated, based on a C_3_/C_4_ competition model that uses the P model to predict C_3_ and C_4_ GPP^30^. Area where cropland cover is higher than 50% were excluded in C_4_ vegetation fraction map to avoid anthropogenic influences. All calculations and analyses were conducted in the open-source environments R (version 4.1.2) and Python (version 3.10).

### Environmental data

We downloaded globally averaged monthly mean CO_2_ concentrations (μmol mol^−1^) from the NOAA Global Monitoring Laboratory for 2000–2017 (NOAA/GML; https://gml.noaa.gov/ccgg/trends/; last access June 2022). Monthly precipitation, maximum, minimum and mean temperature and water vapour pressure at 0.5° resolution were derived from the Climate Research Unit (CRU) TS4.04 data set^31^. Vapour pressure deficit was calculated using maximum and minimum temperatures and water vapour pressure. Atmospheric pressure was estimated from global gridded elevation at 0.5°resolution in the WFDEI meteorological forcing dataset^32^. Hourly surface downwelling shortwave radiation was downloaded from the WFDE5 dataset version 2.0 (ref. 33) and summed to provide monthly totals.

The CRU and WFDE5 data sets have relatively low spatial resolution and are consequently less suitable for site-level model evaluation. We obtained high-resolution climate data from the Climatologies at High resolution for the Earth’s Land Surface Areas (CHELSA) dataset^34^ and used this as the climate forcing at the site scale. The CHELSA data set provides monthly air temperature, precipitation, solar radiation and vapour pressure deficit with a spatial resolution of 30 arc seconds (around 1km) ^34^.

Root-zone soil moisture from the Global Land Evaporation Amsterdam Model (GLEAM) v3.6a product^35,36^ available for the period 1980-2021 was used to estimate soil water stress on the stomatal limitation of photosynthesis.

Annual tree cover percentages for 2000–2020 were derived from MODIS MOD44B v006 (ref. 37). Cropland cover at 0.05°resolution from 2001 –2016 was derived from MODIS MCD12C1 v006 (ref. 38) and regridded to 0.5°resolution using the first order conservative remapping function (*remapcon*) from the Climate Data Operators (CDO) software package (https://code.mpimet.mpg.de/projects/cdo).

### Prediction of annual maximum fAPAR

We hypothesize that on annual and longer time scales, the allocation of carbon to foliage is limited either by water supply, as a transpiring canopy cannot be sustained if insufficient root-zone water leads to prolonged stomatal closure; or by photosynthesis, as building and maintaining leaves (and supplying them with water and nutrients) implies a carbon cost that cannot for long exceed the rate at which they fix carbon. We refer to these two cases as ‘water limited’ and ‘energy limited’. Under water limitation, we hypothesize that plants collectively adjust their rooting behaviour to extract a fraction of annual precipitation from the soil, regardless of its distribution through the year, and allocate carbon to leaves in such a way that all of this water is transpired and GPP consequently maximized. Under energy limitation, we hypothesize that plants allocate carbon to leaves in such a way as to maximize GPP after subtracting the costs of constructing and maintaining leaves and keeping them supplied with water and nutrients. This criterion leads to a well-defined optimum because investment in leaf tissue produces a diminishing return, due to the mutual shading of leaves.

General expressions for GPP and transpiration are:

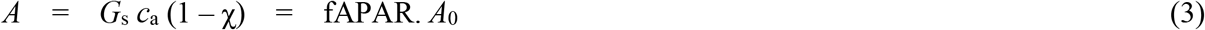

and

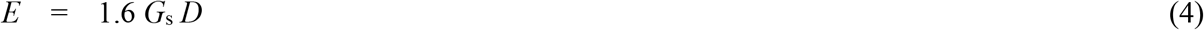

where *A* is GPP (we neglect leaf respiration for simplicity), *E* is transpiration, and *G*_s_ is the canopy conductance for CO_2_. If we also assume that *E* = *f*_0_. *P* then re-arrangement immediately yields equation (1) as the water-limited fAPAR. We now define the net carbon profit (*P*_n_) as:

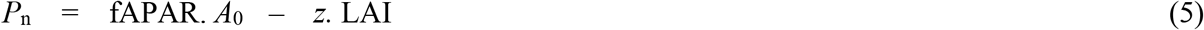

and assume Beer’s law:

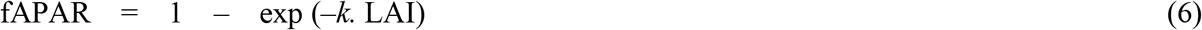

hence

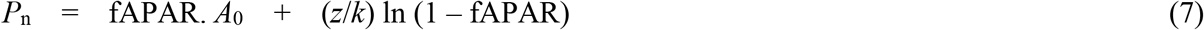

and

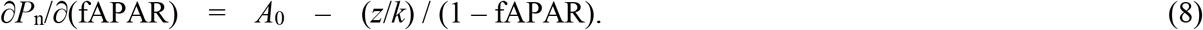

Setting equation (8) to zero yields equation (2), corresponding to a maximum in *P*_n_. The globally fitted values for *z* and *f*_0_ were 13.86 mol m^−2^ a^−1^ and 0.62 (unitless) respectively, obtained by non-linear least regression using the logsumexp function to approximate the minimum function.

### Evaluation of predicted annual maximum fAPAR and its trend

We evaluated the modelled global fAPAR using the 0.05°resolution fAPAR data^39^ at daily timestep from 2000-2017 derived from MODIS MOD15A2H Leaf Area Index/FPAR product^40^. The annual maximum fAPAR were derived first by selecting the maximum value through the year, and then the 95^th^ percentile of each 10×10 0.05°grid cells was used as the maximum fAPAR for each 0.5°grid cell.

To compare the performance of our framework with state-of-the-art dynamic global vegetation models, we downloaded LAI over 2000–2017 as simulated by the ensemble of 15 TRENDY^41^ ecosystem models (TRENDY v9; Table 1) (note that the models do not report fAPAR and generally do not model it separately from LAI). We used the S2 simulations, in which identical, time-varying climate and CO_2_ are prescribed to all the models. TRENDY annual maximum LAI values were converted to fAPAR_max_ using equation (6) for comparison with modelled fAPAR_max_.

**Table 1:** Details of the models from TRENDY v9

| Model name | Spatial resolution | Reference |
| --- | --- | --- |
| CABLE-POP | $1^\circ \times 1^\circ$ | Haverd et al. (2018), ref 42 |
| CLASSIC | $2.8125^\circ \times 2.8125^\circ$ | Melton et al. (2020), ref 43 |
| CLM 5.0 | $0.9375^\circ \times 1.25^\circ$ | Lawrence et al. (2019), ref 44 |
| IBIS | $1^\circ \times 1^\circ$ | Yuan et al. (2014), ref 45 |
| ISAM | $0.5^\circ \times 0.5^\circ$ | Meiyappan et al. (2015), ref 46 |
| ISBA-CTRIP | $1^\circ \times 1^\circ$ | Delire et al. (2020), ref 47 |
| JSBACH | $1.875^\circ \times 1.875^\circ$ | Reick et al. (2021), ref 48 |
| JULES-ES | $1.25^\circ \times 1.875^\circ$ | Wiltshire et al. (2021), ref 49 |
| LPJ-GUESS | $0.5^\circ \times 0.5^\circ$ | Smith et al. (2014), ref 50 |
| LPX-Bern | $0.5^\circ \times 0.5^\circ$ | Lienert and Joos (2018), ref 51 |
| OCN | $1^\circ \times 1^\circ$ | Zaehle and Friend (2010), ref 52 |
| ORCHIDEEv3 | $0.5^\circ \times 0.5^\circ$ | Vuichard et al. (2017), ref 53 |
| SDGVM | $1^\circ \times 1^\circ$ | Walker et al. (2017), ref 54 |
| VISIT | $0.5^\circ \times 0.5^\circ$ | Kato et al. (2013), ref 55 |
| YIBs | $1^\circ \times 1^\circ$ | Yue and Unger (2015), ref 56 |

We visually compared the multi-year average of modelled and observed fAPAR during 2000-2017 across the globe and evaluated the predicted fAPAR_max_ using R-squared (*R*^2^) and root mean squared error (RMSE). We also performed ordinary least-squares linear regression of observed versus predicted fAPAR_max_.

fAPAR trends over the study period (2000–2017) were quantified using Mann-Kendall coefficients at a coarser resolution of 1.5° in order to reduce the incidence of false (positive or negative) trends. Areas with high fAPAR_max_ values (> 0.85) in the initial year were excluded from the analysis. Besides MODIS fAPAR data, we also considered remotely-sensed greenness trends over the same period in the GEOV2^57^, GLOBMAP^58^ and AVHRR^59^ products. We retained only those grid cells where either three or all four products showed trends with the same sign. Predicted (MODIS) and observed trend coefficients were then compared in the same way as the predicted and observed fAPAR_max_.

In-situ measurements provided additional evaluation of model performance. A total of 222 site-year LAI measurements were obtained from the Ground-Based Observations for Validation (GBOV) of Copernicus Global Land Products (https://gbov.acri.fr) and the OLIVE ground database^60,15^. fAPAR was estimated from LAI with equation (6). We determined the annual maximum fAPAR as the fAPAR at peak greenness, and evaluated the predicted against observed fAPAR_max_ at the ground sites using *R*^2^ and RMSE.

### Determination of dominant drivers of the modelled trends in fAPAR

To investigate the causes of fAPAR trends during the study period we conducted simulations in which one of the environmental variables (temperature, solar radiation, precipitation, soil moisture and CO_2_ level) at a time was held constant at its initial-year value, while other variables were allowed to change. The trends in fAPAR_max_ derived from these experiments at each grid cell were compared to the trends seen in the original simulation with all factors varying. The variable that produced the highest reduction in the trend was considered to be the factor controlling the trend. We also examined if this factor had a positive or negative impact on the modelled fAPAR_max_ trend.

## Acknowledgments

WC is funded by the Chinese Scholarship Council. ICP acknowledges support from the European Research Council under the European Union’s Horizon 2020 research and innovation programme (Grant Agreement No: 787203 REALM). ZZ and WH acknowledge support from the National Natural Science Foundation of China (32022052, 91837312, 31971495) and the Tsinghua University Initiative Scientific Research Program (20223080041). SPH acknowledges support from the ERC-funded project GC2.0 (Global Change 2.0: Unlocking the past for a clearer future, No. 694481). This research is a contribution to the Land Ecosystem Models based On New Theory, obseRvations and ExperimEnts (LEMONTREE) project funded through the generosity of Eric and Wendy Schmidt by recommendation of the Schmidt Futures program. All authors acknowledge support from LEMONTREE.

## Author Contributions

ICP, SPH and WH conceived the research; WC and ZZ carried out the data processing and analyses and produced the graphics; all authors contributed to the interpretation of the results. WC produced the first complete draft, with input from ICP and ZZ. All authors contributed to the final version.

## Competing interests

The authors declare that they have no competing interests.

## Notes

### Competing Interest Statement

The authors have declared no competing interest.

